# Effectiveness of direct alignment between ^18^F-Fluorodeoxyglucose images and a standard-space template

**DOI:** 10.1101/2025.09.12.675968

**Authors:** Elizaveta Baranova, Niall W. Duncan

**Affiliations:** Graduate Institute of Mind, Brain and Consciousness, Taipei Medical University, Taipei, Taiwan

**Keywords:** Normalisation, pre-processing, neuroimaging, group analysis, Alzheimer’s disease, glucose

## Abstract

Group analyses in neuroimaging often require spatial normalisation. The most common approach for this involves the use of anatomical images as intermediate references. A lack of a usable anatomical image may then lead to a participant’s data being excluded. This exclusion is a particular concern for challenging patient populations, such as those with movement disorders, and can introduce bias. We evaluated a direct alignment approach for ^1^8F-Fluorodeoxyglucose (FDG-PET) images to a simulated PET standard-space template, which would eliminate the need for an anatomical scan. The performance of this method was compared against the conventional T1-weighted image-based alignment using data from 40 participants with Alzheimer’s disease and 47 healthy controls. Our results show that the direct alignment method performed as well as the conventional method in terms of image-template similarity. It also produced comparable region-of-interest (ROI) metabolic activity values and yielded more extensive and statistically significant group differences in a voxel-wise comparison of Alzheimer’s patients versus healthy controls. The findings suggest that direct alignment is a robust and effective alternative for FDG-PET image normalisation.

## Introduction

Group analyses of neuroimaging data generally require that the images from each participant be put into a shared standard space to allow voxel-wise or region-of-interest comparisons across individuals. This will often require an indirect, multi-step process, where the data of interest is first aligned with a separate anatomical image and then registered to a standard template via this image. In the case of PET imaging, this anatomical image can be a CT scan acquired on the PET scanner itself. Alternatively, an MRI image may be acquired separately or, as may become more common, on a combined PET-MRI scanner (9).

With this spatial normalisation step being necessary in most cases, PET data from participants for whom anatomical images are missing or unusable may need to be excluded from an analysis. In the case of MRI, this may be due to poor quality images or from participant status preventing or interrupting scans. Such occurrences can be common in challenging patient groups, such as those who display high head motion, people with metal implants, or groups that have difficulty with compliance with MRI procedures. With CT, is-sues of image quality can also arise through head motion or technical factors (8), as can misalignments between CT and PET images (11, 12). CT-based alignment also faces challenges due to the limited neuroanatomical information visible through this method.

The exclusion of PET datasets for reasons of missing or unusable anatomical images is wasteful. Systematic exclusion could also introduce biases by disproportionately removing data from patient populations with greater movement disorders or cognitive impairments, thereby limiting the accuracy and generalisability of findings. As such, it would be advantageous to have a robust method for image normalisation that relies on the PET image alone. This would mean that all participants with usable PET data can be included in an analysis, independent of the need for anatomical images.

Similar questions around image co-registration exist for other neuroimaging modalities, such as functional MRI (fMRI). In the fMRI context, direct template alignment has been suggested as an alternative approach to anatomical image-based registration (2, 4, 5). For this, a specific EPI template image is used, rather than the more common T1-weighted anatomical image, so that the template image properties are matched with the images to be aligned to it. This matching of image properties with those to be aligned allows optimal performance of the registration algorithms and can allow the alignment to account for the specific features of the data. Such direct alignment can be with either a standard-space template (2), such as the MNI152 template, or with a study-specific template (4, 5). In the case of EPI images, this type of direct standard-space template alignment can give superior outcomes to the more common indirect methods (2, 4).

In this work, we extend the direct alignment approach to the case of ^18^F-Fluorodeoxyglucose PET (FDG-PET). The objective was to establish a spatial normalisation approach that would allow accurate group analysis of this modality, independent of the need to have a usable anatomical image. In particular, we focus on the alignment of FDG-PET images from a group of participants to a standard-space template. The effectiveness of such a direct approach was then compared to the conventional multi-stage approach using the participant’s T1-weighted anatomical image.

## Methods

### Dataset

Data from the Alzheimer’s Disease Neuroimaging Initiative (ADNI) were used. From this, we selected people between 50 and 70 years of age who had both a T1-weighted anatomical scan (MPRAGE or FSPGR) and PET scans with an 18F-FDG radiotracer. This gave 112 participants. This group was a mix of people with Alzheimer’s disease (AD) and healthy controls (HC). These were further filtered to exclude those that had data in a format other than NIfTI (12 participants) or who did not have dynamic data in six five minute bins (12 participants). An additional participant was excluded as they had an abnormal anatomical feature. The final sample of 87 people included 40 with Alzheimer’s disease and 47 controls (mean age = 65.5 ± 3.72 years, age range = 55-69, 36 females). Individual details are given in Supplementary table 1.

### Anatomical image preprocessing

T1-weighted anatomical images underwent the following processing steps: 1) Reduction of the field of view in the inferior-superior axis to exclude excess neck tissue (*robustfov*); 2) bias-field correction with an N4 algorithm implemented in ANTs (16); segmentation of the image through FastSurfer (6, 7); application of the brain mask calculated by FastSurfer to remove non-brain tissue. Anatomical images were kept in their native resolution (see Supplementary table 1 for details).

### FDG-PET image preprocessing

Arterial sampling or images sufficient to create an artery reference region were not available and so the dynamic frames were combined by calculating the temporal mean. Although not optimal for analysis of the underlying metabolic properties, this approach produces an image sufficient for the aim of testing image alignment quality as image values are not to be compared between individuals. Mean FDG-PET images were then skull-stripped with FSL’s BET tool. Data were acquired at different sites with different scanners and so FDG-PET image resolutions differed between participants, ranging from 1.02 x 1.02 x 2.00 mm^3^ to 2.00 x 2.00 x 4.25 mm^3^ (see Supplementary table 1 for details).

### Template creation

We sought to create a template image that had the highest intensity in grey-matter, with lower intensity non-brain tissue areas. White-matter and cerebrospinal fluid areas were left empty to provide a clear tissue boundary. To create this simulated image, we first segmented skull-stripped T1-weighted MNI152NLin6Asym images, as included with FSL, into grey-matter, white-matter, and cerebrospinal fluid with FSL’s *FAST*. This was done for both the 1 mm and 2 mm versions to allow comparison between different template base resolutions. Grey matter proportion maps were then convolved with a Gaussian kernel to simulate the dispersal seen in FDG-PET images. The question of which size of kernel to apply will be returned to below. Each voxel was then multiplied by a constant value so that their overall intensities were higher than non-brain tissue. The brain was then removed from the full T1-weighted MNI152 images and the simulated brain map inserted. An example of a simulated template image is shown in Figure 1A.

**Figure 1.**
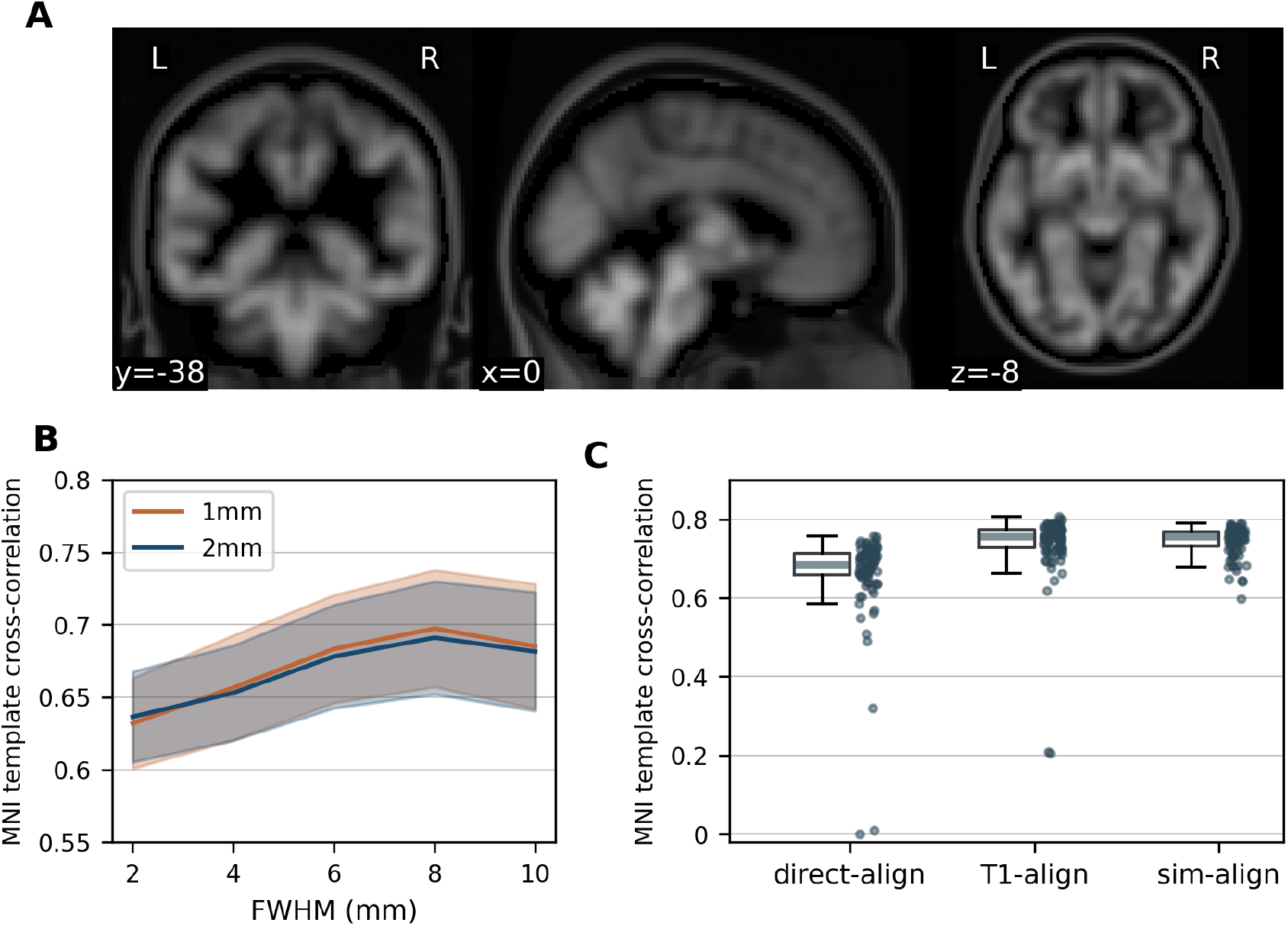
**A**. Example simulated PET template (8 mm FWHM smoothing). **B**. Similarity between participant images aligned to standard space via simulated PET templates with different smoothing levels and the MNI template image. Analysis was done on a subset of 10 participants with templates at 1 mm and 2 mm resolution. Shading denotes standard deviation across participants. **C**. Similarity between all participant images in MNI space and the MNI template for different template types.

### FDG-PET to template alignment

Three different alignment approaches were compared. Each alignment was calculated with the ANTs *antsRegister* tool (1). In each case, a rigid transform was first calculated to bring the images into the same space. This was followed by a linear registration and, finally, a non-linear SyN registration. No mask was used for the rigid and linear registrations so that large-scale information such as skull position could be exploited. A mask of just the cortex, midbrain, and cerebellum was used for the final, non-linear, registration. Registrations were calculated with images that included non-brain tissue. The transformations were then applied to skull-stripped FDG-PET images with the *antsApplyTransforms* tool.

The most basic registration approach involved aligning the mean FDG-PET images directly to the T1-weighted MNI152 template (*basic-align*). This method was assumed to be the least effective and was taken as a comparator for the other two, more complex, approaches. The second approach was a standard multi-step alignment via participant’s T1-weighted anatomical image (*T1-align*). The third ap-proach directly aligned participant FDG-PET images to the simulated FDG-PET image in MNI152 space (*sim-align*).

### Base resolution and kernel size

To establish what base resolution (1 mm or 2 mm) and kernel size was optimal, we selected ten participants with each of the various FDG-PET resolutions and calculated sim-align registrations against simulated PET templates created with different kernels. These varied from 2 mm to 10 mm, in steps of 2 mm. The cross-correlation between the aligned images and the T1-weighted MNI template was calculated for each and then plotted.

### Comparison between methods - PET-template similarity

A set of different approaches were taken to compare the effectiveness of each registration method. Firstly, the cross-correlation between the normalised PET images and the MNI template was calculated for each participant. This metric was limited to within the cortex, the midbrain and the cerebellum, with non-brain tissue excluded. Correlation coefficients were then compared between the methods with a non-parametric Friedman test, followed by Nemenyi post-hoc tests. The group variability of alignment quality was calculated as the median absolute deviation of these correlation coefficients. These were statistically compared between methods through a bootstrapping approach. Statistical analyses were carried out with *numpy* and *scipy* Python packages. Figures were created with the *matplotlib* and *nilearn* packages.

### Comparison between methods - ROI similarity

Next, mean normalised FDG-PET values for each cortical and subcortical region of interest (ROI) from the Harvard-Oxford atlas were calculated for each subject. Participant images were normalised by dividing values within the brain by the trimmed mean (5th to 95th percentile) of non-brain tissue intensity values (14). Non-brain tissue was defined as the areas outside the brain in the MNI152 template and included the face, skull, and neck. These values were then averaged within each ROI and correlated between the *T1-align* and *sim-align* data (Spearman’s correlation) to establish the degree of similarity between these methods for ROI-based analysis.

### Comparison between methods - Voxel-wise group difference

Next, normalised FDG-PET images in template space from patients with Alzheimer’s were compared to those from healthy controls in a voxel-wise comparison of cortical grey-matter. Images were downsampled to 3 mm isotropic prior to this analysis. The comparison was carried out with FSL’s randomise tool, using 10,000 permutations (17). Voxel-wise family-wise error rates were controlled at *α* = 0.01.

## Results

### Template optimisation

Testing different smoothing kernels when creating the simulated PET template found that the optimal kernel was one with a FWHM of 8 mm. The alignment performance was equivalent for both the 1 mm and 2 mm resolution templates (Figure 1B). As such, the simulated PET template created at 2 mm resolution with 8 mm smoothing was used for subsequent analyses.

### PET-template similarity

PET-to-template cross-correlation differed between the three methods 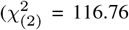, *p* = 4.43e-26; Figure 1C). Post-hoc tests showed that the *sim-align* (*M* = 0.74) method performed as well as the *T1-align* (*M* = 0.74, *p* = 0.63) and better than the *basic-align* (*M* = 0.66, *p* = 0.0) methods. The *T1-align* method also performed better than *basic-align* (*p* = 0.0). Variation in alignment quality across participants was equivalent for the *sim-align* (*MAD* = 0.02, 95% CI = 0.01 - 0.02) and *T1-align* methods (*MAD* = 0.02, 95% CI = 0.02 - 0.03, *p* = 0.084). Variation for the *sim-align* method was lower than that for *basic-align* (*MAD* = 0.03, 95% CI = 0.02 - 0.04, *p* = 4.0e-4).

Alignment quality with the *sim-align* method was not related to the resolution of the participant’s PET image (sagittal: *r* = 0.13, *p* = 0.23; axial: *r* = 0.07, *p* = 0.52).

## ROI similarity

The median Spearman’s correlation between *sim-align* and *T1-align* images for normalised ROI FDG-PET values was 0.85 (range = 0.77-0.93). The median correlation in cortical regions was 0.86 (range = 0.79-0.93), with subcortical regions showing a median correlation of 0.84 (range = 0.77-0.86). The highest similarity was observed in the temporooccipital inferior temporal gyrus (*r* = 0.93) and the lowest in the left putamen (*r* = 0.77). Regional correlation values are shown in Figure 2 and Supplementary table 2.

**Figure 2.**
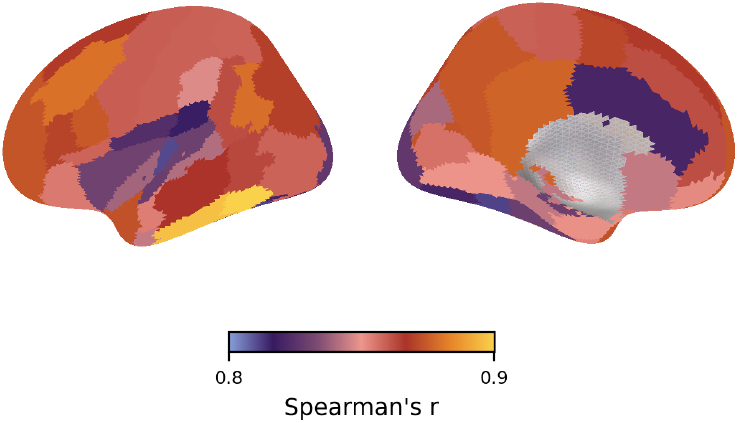
Regional Spearman’s correlation values between PET images aligned to standard space with *T1-align* and *sim-align* methods. (Subcortical regions are not shown.)

### Voxel-wise group difference

As shown in Figure 3, voxel-wise analysis with images aligned to standard space via the *sim-align* method showed more extensive group differences at a voxel-wise FWE corrected threshold of *α* = 0.01 than did those using the *T1-align* method (5109 voxels compared to 2258). The peak contrast statistical value was also higher for *sim-align* images (*T* = 8.89) than *t1-align* (*T* = 7.49). Unthresholded images are shown in Supplementary figure 1.

**Figure 3.**
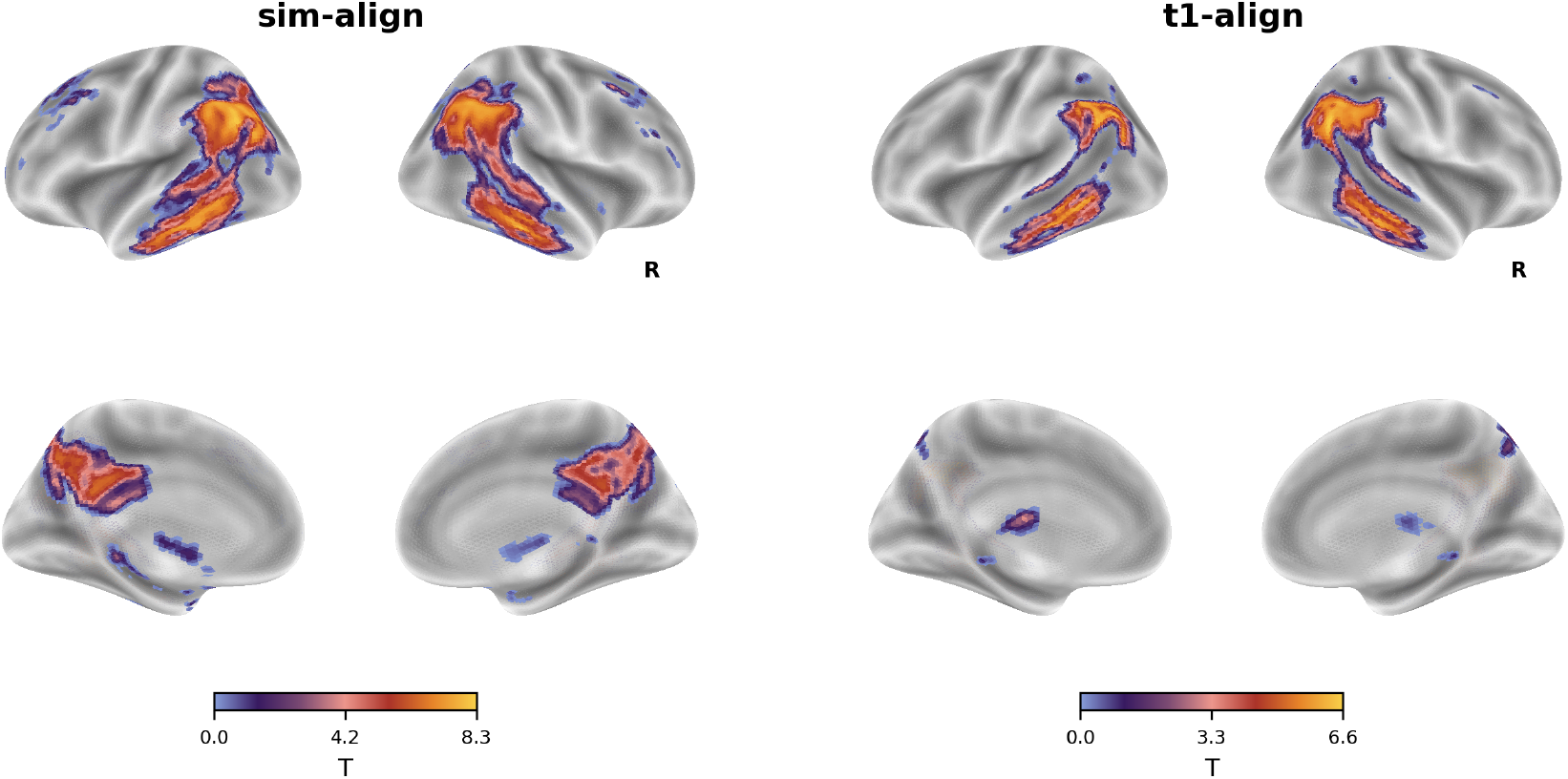
Contrast images for comparison of HC with AD. Images are thresholded at a voxel-wise FWE corrected *α* = 0.01.

## Discussion

Direct alignment between FDG-PET images and a simulated PET template was found to perform as well as alignment via an individual T1-weighted image. This direct alignment technique was found to produce similar ROI metabolic activity estimates to the T1-based alignment approach. It was also found to give improved outcomes in a voxel-wise comparison between healthy controls and patients with Alzheimer’s disease. This level of performance echoes that seen with direct alignment methods for EPI data (2, 4, 5).

The use of a direct alignment technique has the potential to reduce the number of scans a participant must undertake, particularly in cases where an additional T1-scan is only being acquired for the purpose of PET normalisation. This could reduce research costs and the burden on participants. The latter point is of particular value in research groups, such as those with particular disorders or people with little free time, where attending multiple scanning sessions can be stressful or inconvenient. At the same time, direct alignment can help solve problems resulting from PET-CT misalignments or CT image corruption. It may also play a role in reducing radiation exposure risk if the number of CT scans required can be reduced (13). Finally, the direct approach has a small advantage in terms of computational complexity over two-stage T1-based alignment.

The MNI152NLin6Asym template was used here, but the creation of a simulated PET template would in principle be possible with any standard space template where grey matter distribution can be established. Likewise, study-specific templates could be produced in situations where this is required. Conversion of data to a surface, rather than a volume, for subsequent analysis would be possible through standard means (4). Once in a standard space, different atlases are easily applied for data analysis and the data becomes more amenable for sharing and future mega-analysis (3, 10).

The effectiveness of direct alignment for FDG-PET images is demonstrated here but there is scope for this approach to be applied to different radiotracers. This will depend upon the availability of a suitable reference substrate to base the simulated image on (grey matter density in this case). An interesting potential approach for some tracers may be to use gene expression patterns for target receptors or transporters, such as those available through the Allen Human Brain Atlas (15). For example, expression patterns for the dopamine transporter gene may form an effective base for a simulated PET template for different radiotracers for this protein. Testing such an approach may be a useful future research direction.

A potential limitation of our analysis here is that it was done with images from older participants, including those with pathological cortical atrophy, due to data availability. These populations are generally more difficult to achieve good quality alignments to standard space templates and so the good performance here may translate to good performance in less challenging groups. Some testing in specific applications may be warranted, however. The dataset used here also consisted of a mixture of spatial resolutions and scanner types. Although this may be seen as an advantage in terms of demonstrating the robustness of the technique, it did mean that we aimed for optimising overall performance across all data. Where a study only has one resolution, some testing of parameters for study-specific performance may further improve outcomes.

In summary, we demonstrate the effectiveness of a direct alignment approach for FDG-PET images. This approach could have a variety of advantages. Overall, the approach appears to merit consideration as part of FDG-PET analysis pipelines.

## Data and code availability

ADNI data is available at https://adni.loni.usc.edu/data-samples/adni-data/. Data and code required to reproduce the results presented here are available at https://osf.io/egy6x/.

## Acknowledgments

This work was supported by grants from the Taiwan Ministry of Science and Technology to NWD (NSTC113-2423-H-038-002-MY3; NSTC110-2628-H-038-001-MY4). The preprint was created using a modified version of Ricardo Henriques’ BioRxiv LaTex template (https://henriqueslab.github.io/resources/bioRxivTemplate/).

## Contributions

EB: Conceptualization, Methodology, Formal Analysis, Writing – original draft

NWD: Conceptualization, Methodology, Formal Analysis, Visualization, Writing – original draft

## Supplementary materials

**Table 1.** Information for each participant. F = female; M = male, AD = person with Alzheimer’s disease; HC = healthy control

| Participant | Age (years) | Sex | Group | T1 voxel size (mm<sup>3</sup>) | PET voxel size (mm<sup>3</sup>) |
| --- | --- | --- | --- | --- | --- |
| 1 | 66.8 | M | AD | 1.20 x 1.02 x 1.02 | 1.02 x 1.02 x 2.03 |
| 2 | 60.8 | M | AD | 1.20 x 1.02 x 1.02 | 1.02 x 1.02 x 2.03 |
| 3 | 68.8 | F | HC | 1.20 x 1.02 x 1.02 | 1.02 x 1.02 x 2.03 |
| 4 | 65.9 | F | HC | 1.20 x 1.02 x 1.02 | 1.02 x 1.02 x 2.03 |
| 5 | 67.7 | F | HC | 1.20 x 1.02 x 1.02 | 1.02 x 1.02 x 2.03 |
| 6 | 68.6 | F | HC | 1.20 x 1.02 x 1.02 | 1.02 x 1.02 x 2.03 |
| 7 | 59.9 | F | HC | 1.20 x 1.02 x 1.02 | 1.02 x 1.02 x 2.03 |
| 8 | 61.5 | F | AD | 1.20 x 1.02 x 1.02 | 1.02 x 1.02 x 2.00 |
| 9 | 55.3 | M | AD | 1.00 x 1.00 x 1.00 | 1.02 x 1.02 x 2.03 |
| 10 | 67.5 | M | AD | 1.20 x 0.94 x 0.94 | 2.06 x 2.06 x 2.43 |
| 11 | 68.0 | M | AD | 1.20 x 1.02 x 1.02 | 2.57 x 2.57 x 2.43 |
| 12 | 67.0 | F | HC | 1.20 x 1.00 x 1.00 | 2.57 x 2.57 x 2.43 |
| 13 | 59.0 | F | AD | 1.20 x 0.94 x 0.94 | 2.06 x 2.06 x 3.38 |
| 14 | 66.9 | M | HC | 1.20 x 1.00 x 1.00 | 2.00 x 2.00 x 4.25 |
| 15 | 68.5 | F | HC | 1.20 x 1.00 x 1.00 | 2.00 x 2.00 x 4.25 |
| 16 | 67.5 | M | AD | 1.20 x 1.00 x 1.00 | 2.00 x 2.00 x 4.25 |
| 17 | 56.6 | M | AD | 1.20 x 1.00 x 1.00 | 2.00 x 2.00 x 4.25 |
| 18 | 65.5 | M | HC | 1.20 x 1.25 x 1.25 | 2.06 x 2.06 x 2.43 |
| 19 | 68.0 | F | AD | 1.20 x 1.00 x 1.00 | 1.02 x 1.02 x 2.03 |
| 20 | 69.2 | F | AD | 1.20 x 1.00 x 1.00 | 1.02 x 1.02 x 2.03 |
| 21 | 67.1 | F | HC | 1.20 x 1.00 x 1.00 | 2.00 x 2.00 x 4.25 |
| 22 | 65.2 | F | HC | 1.20 x 1.00 x 1.00 | 2.00 x 2.00 x 4.25 |
| 23 | 69.8 | F | HC | 1.20 x 1.00 x 1.00 | 2.00 x 2.00 x 3.27 |
| 24 | 56.0 | M | AD | 1.20 x 1.00 x 1.00 | 2.00 x 2.00 x 2.00 |
| 25 | 67.5 | F | HC | 1.20 x 1.00 x 1.00 | 2.00 x 2.00 x 2.00 |
| 26 | 55.7 | F | AD | 1.20 x 1.02 x 1.02 | 2.03 x 2.03 x 2.00 |
| 27 | 66.0 | F | AD | 1.20 x 1.02 x 1.02 | 2.03 x 2.03 x 2.00 |
| 28 | 69.5 | F | HC | 1.20 x 1.02 x 1.02 | 1.02 x 1.02 x 2.00 |
| 29 | 66.3 | F | AD | 1.20 x 1.02 x 1.02 | 1.02 x 1.02 x 2.00 |
| 30 | 63.0 | F | AD | 1.20 x 1.00 x 1.00 | 2.00 x 2.00 x 3.27 |
| 31 | 68.5 | F | HC | 1.20 x 1.00 x 1.00 | 2.57 x 2.57 x 2.43 |
| 32 | 69.4 | F | HC | 1.20 x 1.05 x 1.05 | 2.00 x 2.00 x 2.00 |
| 33 | 65.8 | F | HC | 1.20 x 1.02 x 1.02 | 2.00 x 2.00 x 2.00 |
| 34 | 60.5 | M | AD | 1.20 x 0.94 x 0.94 | 2.00 x 2.00 x 3.27 |
| 35 | 68.5 | F | AD | 1.20 x 1.00 x 1.00 | 2.00 x 2.00 x 3.27 |
| 36 | 64.7 | F | AD | 1.20 x 1.00 x 1.00 | 2.00 x 2.00 x 3.27 |
| 37 | 69.3 | F | HC | 1.20 x 1.00 x 1.00 | 2.00 x 2.00 x 3.27 |
| 38 | 62.2 | F | HC | 1.20 x 1.25 x 1.25 | 2.06 x 2.06 x 2.43 |
| 39 | 69.5 | F | AD | 1.20 x 1.25 x 1.25 | 2.06 x 2.06 x 2.43 |
| 40 | 69.4 | M | AD | 1.20 x 1.30 x 1.30 | 2.06 x 2.06 x 2.43 |
| 41 | 60.8 | F | AD | 1.20 x 1.00 x 1.00 | 2.57 x 2.57 x 2.43 |
| 42 | 63.5 | F | HC | 1.20 x 1.00 x 1.00 | 2.57 x 2.57 x 2.43 |
| 43 | 69.2 | F | HC | 1.20 x 1.00 x 1.00 | 2.57 x 2.57 x 2.43 |
| 44 | 68.6 | F | HC | 1.20 x 1.00 x 1.00 | 1.02 x 1.02 x 2.03 |
| 45 | 69.4 | M | AD | 1.20 x 1.00 x 1.00 | 1.02 x 1.02 x 2.03 |
| 46 | 67.6 | F | HC | 1.20 x 1.00 x 1.00 | 1.02 x 1.02 x 2.03 |
| 47 | 68.2 | M | HC | 1.20 x 1.00 x 1.00 | 2.57 x 2.57 x 2.43 |
| 48 | 59.4 | F | AD | 1.20 x 1.00 x 1.00 | 1.02 x 1.02 x 2.03 |
| 49 | 66.8 | F | HC | 1.20 x 1.00 x 1.00 | 2.00 x 2.00 x 3.27 |
| 50 | 66.2 | F | HC | 1.20 x 1.00 x 1.00 | 2.00 x 2.00 x 3.27 |
| 51 | 65.1 | F | HC | 1.20 x 1.00 x 1.00 | 2.00 x 2.00 x 4.25 |
| 52 | 69.9 | M | HC | 1.20 x 1.00 x 1.00 | 2.00 x 2.00 x 3.27 |
| 53 | 63.2 | M | HC | 1.20 x 1.00 x 1.00 | 2.00 x 2.00 x 3.27 |
| 54 | 67.5 | F | HC | 1.20 x 1.00 x 1.00 | 2.00 x 2.00 x 3.27 |
| 55 | 65.2 | M | HC | 1.20 x 1.00 x 1.00 | 2.00 x 2.00 x 3.27 |
| 56 | 61.3 | M | HC | 1.20 x 1.00 x 1.00 | 2.00 x 2.00 x 3.27 |
| 57 | 64.4 | F | AD | 1.20 x 1.00 x 1.00 | 2.00 x 2.00 x 3.27 |
| 58 | 63.6 | F | HC | 1.20 x 1.00 x 1.00 | 2.00 x 2.00 x 3.27 |

Table 1 – continued from previous page
| Participant | Age (years) | Sex | Group | T1 voxel size (mm<sup>3</sup>) | PET voxel size (mm<sup>3</sup>) |
| --- | --- | --- | --- | --- | --- |
| 59 | 66.7 | M | HC | 1.20 x 1.00 x 1.00 | 2.00 x 2.00 x 3.27 |
| 60 | 69.6 | F | AD | 1.20 x 0.94 x 0.94 | 2.57 x 2.57 x 2.43 |
| 61 | 55.2 | F | AD | 1.20 x 0.94 x 0.94 | 2.57 x 2.57 x 2.43 |
| 62 | 69.5 | M | AD | 1.20 x 0.94 x 0.94 | 2.57 x 2.57 x 2.43 |
| 63 | 68.2 | F | HC | 1.20 x 1.02 x 1.02 | 2.57 x 2.57 x 2.43 |
| 64 | 65.6 | M | HC | 1.20 x 1.02 x 1.02 | 2.57 x 2.57 x 2.43 |
| 65 | 64.1 | F | AD | 1.20 x 1.02 x 1.02 | 2.00 x 2.00 x 3.27 |
| 66 | 68.1 | M | HC | 1.20 x 1.00 x 1.00 | 2.00 x 2.00 x 3.27 |
| 67 | 62.3 | M | AD | 1.20 x 1.00 x 1.00 | 2.00 x 2.00 x 3.27 |
| 68 | 66.0 | M | HC | 1.20 x 1.00 x 1.00 | 2.00 x 2.00 x 3.27 |
| 69 | 69.6 | F | HC | 1.20 x 1.00 x 1.00 | 2.00 x 2.00 x 3.27 |
| 70 | 63.9 | M | AD | 1.20 x 1.00 x 1.00 | 2.00 x 2.00 x 3.27 |
| 71 | 67.7 | F | AD | 1.20 x 0.94 x 0.94 | 2.00 x 2.00 x 4.25 |
| 72 | 64.3 | M | AD | 1.20 x 0.94 x 0.94 | 1.95 x 1.95 x 4.25 |
| 73 | 65.1 | M | HC | 1.20 x 1.02 x 1.02 | 2.57 x 2.57 x 2.43 |
| 74 | 61.9 | M | AD | 1.20 x 1.02 x 1.02 | 2.57 x 2.57 x 2.43 |
| 75 | 61.8 | M | AD | 1.20 x 1.02 x 1.02 | 2.57 x 2.57 x 2.43 |
| 76 | 66.0 | M | AD | 1.20 x 1.02 x 1.02 | 2.57 x 2.57 x 2.43 |
| 77 | 68.4 | M | HC | 1.20 x 1.02 x 1.02 | 2.57 x 2.57 x 2.43 |
| 78 | 67.8 | M | HC | 1.20 x 1.02 x 1.02 | 2.57 x 2.57 x 2.43 |
| 79 | 60.9 | F | AD | 1.20 x 1.00 x 1.00 | 1.02 x 1.02 x 2.03 |
| 80 | 68.4 | F | AD | 1.20 x 1.00 x 1.00 | 1.02 x 1.02 x 2.03 |
| 81 | 68.5 | F | HC | 1.20 x 1.00 x 1.00 | 2.57 x 2.57 x 2.43 |
| 82 | 61.2 | M | AD | 1.20 x 1.00 x 1.00 | 2.57 x 2.57 x 2.43 |

**Table 2.** Regional metabolic index values and Spearman’s correlation between *t1-align* and *sim-align* FDG-PET images. Regions are from the Harvard-Oxford cortical and subcortical atlases.

| Participant | *t1-align* mean | *sim-align* mean | Correlation |
| --- | --- | --- | --- |
| Frontal Pole | 6.84 | 6.26 | 0.89 |
| Insular Cortex | 6.48 | 6.01 | 0.82 |
| Superior Frontal Gyrus | 6.81 | 6.21 | 0.88 |
| Middle Frontal Gyrus | 7.27 | 6.68 | 0.90 |
| Inferior Frontal Gyrus, pars triangularis | 7.21 | 6.70 | 0.88 |
| Inferior Frontal Gyrus, pars opercularis | 7.32 | 6.75 | 0.89 |
| Precentral Gyrus | 7.01 | 6.44 | 0.87 |
| Temporal Pole | 4.79 | 4.29 | 0.89 |
| Superior Temporal Gyrus, anterior division | 6.25 | 5.86 | 0.82 |
| Superior Temporal Gyrus, posterior division | 6.77 | 6.38 | 0.84 |
| Middle Temporal Gyrus, anterior division | 5.86 | 5.52 | 0.86 |
| Middle Temporal Gyrus, posterior division | 6.23 | 5.92 | 0.88 |
| Middle Temporal Gyrus, temporooccipital part | 6.42 | 6.04 | 0.87 |
| Inferior Temporal Gyrus, anterior division | 5.64 | 5.16 | 0.84 |
| Inferior Temporal Gyrus, posterior division | 5.85 | 5.27 | 0.93 |
| Inferior Temporal Gyrus, temporooccipital part | 6.32 | 5.83 | 0.93 |
| Postcentral Gyrus | 6.76 | 6.31 | 0.87 |
| Superior Parietal Lobule | 6.55 | 6.03 | 0.87 |
| Supramarginal Gyrus, anterior division | 6.47 | 6.10 | 0.85 |
| Supramarginal Gyrus, posterior division | 6.63 | 6.16 | 0.87 |
| Angular Gyrus | 6.71 | 6.13 | 0.90 |
| Lateral Occipital Cortex, superior division | 6.60 | 6.13 | 0.88 |
| Lateral Occipital Cortex, inferior division | 6.84 | 6.41 | 0.87 |
| Intracalcarine Cortex | 9.39 | 8.21 | 0.86 |
| Frontal Medial Cortex | 7.36 | 6.55 | 0.85 |
| Juxtapositional Lobule Cortex | 7.22 | 6.54 | 0.88 |
| Subcallosal Cortex | 6.06 | 5.42 | 0.84 |
| Paracingulate Gyrus | 7.43 | 6.61 | 0.87 |
| Cingulate Gyrus, anterior division | 6.56 | 5.97 | 0.80 |
| Cingulate Gyrus, posterior division | 7.79 | 6.75 | 0.89 |
| Precuneous Cortex | 7.75 | 6.89 | 0.89 |

**Table 2 – continued from previous page**
| **Participant** | ***t1-align* mean** | ***sim-align* mean** | **Correlation** |
| --- | --- | --- | --- |
| Cuneal Cortex | 8.01 | 7.11 | 0.84 |
| Frontal Orbital Cortex | 6.79 | 5.95 | 0.86 |
| Parahippocampal Gyrus, anterior division | 4.77 | 3.79 | 0.85 |
| Parahippocampal Gyrus, posterior division | 5.38 | 4.71 | 0.84 |
| Lingual Gyrus | 7.65 | 6.76 | 0.85 |
| Temporal Fusiform Cortex, anterior division | 5.39 | 4.79 | 0.86 |
| Temporal Fusiform Cortex, posterior division | 5.92 | 5.24 | 0.82 |
| Temporal Occipital Fusiform Cortex | 7.00 | 6.19 | 0.79 |
| Occipital Fusiform Gyrus | 7.88 | 6.89 | 0.80 |
| Frontal Operculum Cortex | 7.81 | 6.87 | 0.87 |
| Central Opercular Cortex | 7.20 | 6.44 | 0.81 |
| Parietal Operculum Cortex | 7.40 | 6.65 | 0.80 |
| Planum Polare | 5.76 | 5.04 | 0.83 |
| Heschl's Gyrus | 8.17 | 6.90 | 0.79 |
| Planum Temporale | 7.34 | 6.56 | 0.82 |
| Supracalcarine Cortex | 9.27 | 8.27 | 0.89 |
| Occipital Pole | 7.00 | 6.74 | 0.81 |
| Left Thalamus | 6.47 | 6.09 | 0.84 |
| Left Caudate | 8.73 | 7.06 | 0.84 |
| Left Putamen | 6.07 | 4.67 | 0.77 |
| Left Hippocampus | 4.79 | 4.41 | 0.86 |
| Left Amygdala | 7.66 | 7.35 | 0.84 |
| Right Thalamus | 6.06 | 5.94 | 0.79 |
| Right Caudate | 8.69 | 7.05 | 0.84 |
| Right Putamen | 5.96 | 4.69 | 0.79 |
| Right Hippocampus | 4.76 | 4.36 | 0.84 |
| Right Amygdala | 7.36 | 7.00 | 0.81 |

**Supplementary Fig. 1.**
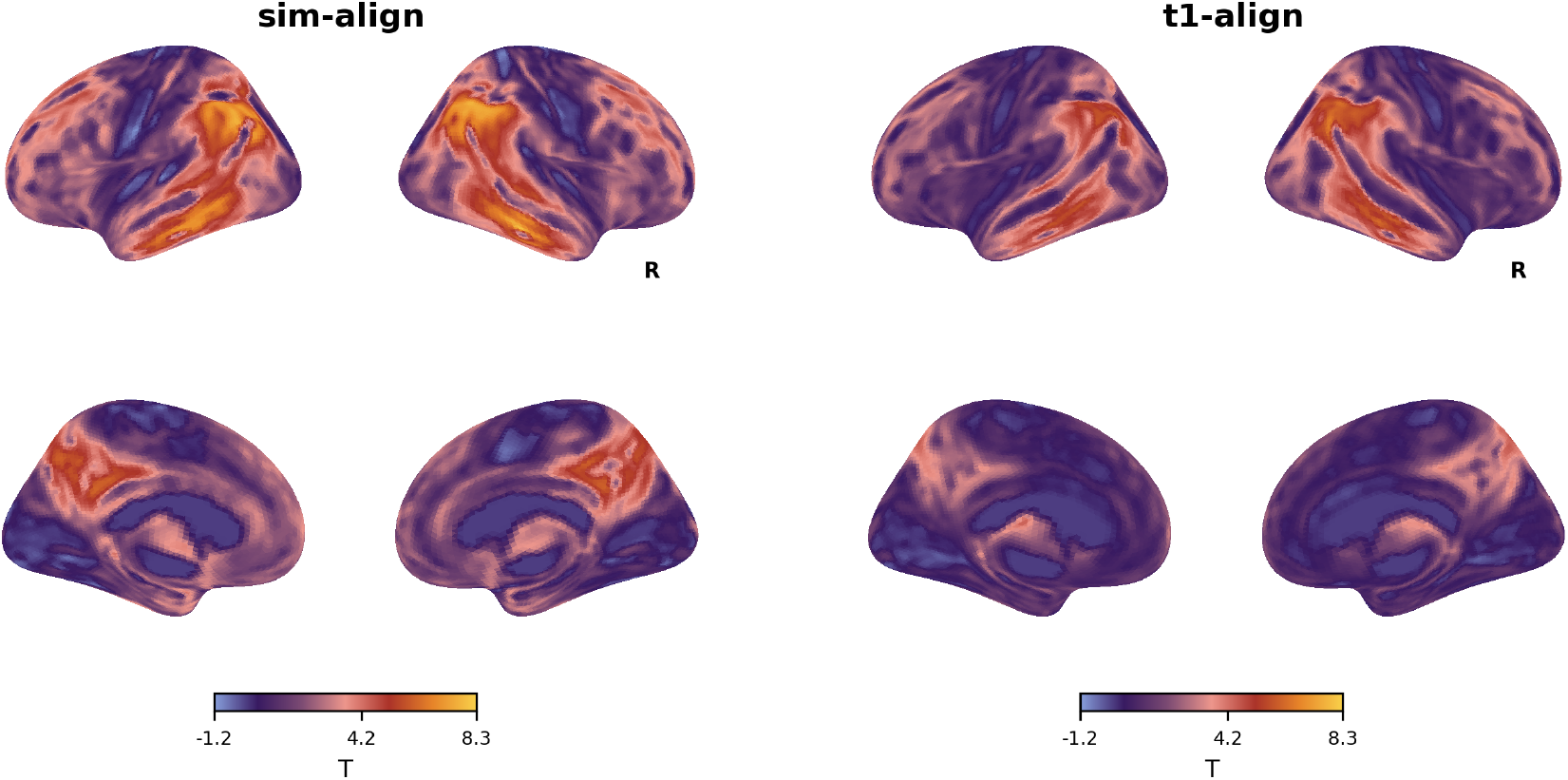
Unthresholded statistical maps for the comparison of healthy controls with patients with Alzheimer’s.

